# Altered Hippocampal Structure-Function Coupling and Brain-Cognition Aging Profile in Mild Cognitive Impairment

**DOI:** 10.64898/2026.06.19.733422

**Authors:** Damien Marie, Dimitra Kokkinou, Chantal Junker-Tschopp, Matthias Kliegel, Gilles Allali, Andrea Brioschi Guevara, Giovanni B. Frisoni, Clara E. James

## Abstract

Hippocampal atrophy and altered functional connectivity are prominent features of mild cognitive impairment (MCI), yet their relationships and joint contribution to cognitive impairment remain unclear. We evaluated multidomain cognitive-sensorimotor performance, whole-brain gray matter atrophy patterns, associated functional connectivity, and their coupling in MCI and HC. We conducted this cross-sectional study in 25 clinically diagnosed amnestic and non-amnestic MCI patients and 15 age- and education-matched healthy controls (HC). Participants (57-81 years old) completed a battery assessing general cognition, executive functions, attention, speech-in-noise perception, manual dexterity, and balance, combined with structural and resting-state functional magnetic resonance imaging. Among the 25 MCI patients, 8 (32%) presented with amnestic MCI, 15 (60%) with non-amnestic MCI, and 2 (8%) with subjective cognitive decline. Linear mixed models revealed that semantic fluency (g = 0.74) was the largest discriminator of MCI, followed by attention (g = 0.61), phonemic fluency (g = 0.59), and speech-in-noise perception (g = 0.56). Voxel-based morphometry showed bilateral hippocampal and anteromedial cerebellar gray matter atrophy in MCI. Seed-based functional connectivity analyses indicated that atrophy was not uniformly associated with reduced connectivity. Cortico-subcortical hypoconnectivity emerged only in networks associated with left hippocampal atrophy in MCI compared with HC. Network-based analyses showed disrupted brain-behavior coupling and a stronger detrimental influence of age in MCI than in HC. These findings confirm the critical role of hippocampal structure-function relationships in multidomain MCI symptoms. A decoupled brain-cognition aging profile suggests that multimodal indices integrating hippocampal structure, connectivity, and behavioral performance may strengthen MCI diagnosis.

## Introduction

Mild Cognitive Impairment (MCI) is a clinical construct characterized by acquired cognitive impairment without significant functional limitation (Frisoni et al., 2023; Petersen et al., 1995). Prevalence (10-15% of older adults ≥ 65 years old) and dementia risk are high (5-20% annual conversion rate) (Anderson, 2019; Langa and Levine, 2014). MCI can affect multiple cognitive domains, including memory, attention, and executive function (Frisoni et al., 2023). Speech-in-noise perception deficits occur (Lee et al., 2016), which may relate to executive dysfunction (Lee et al., 2018). Sensorimotor domains such as gait or balance can also be impaired (Bahureksa et al., 2016), contributing to increased fall risk (Delbaere et al., 2012).

In addition to sensorimotor and cognitive deficits, MCI is associated with multimodal neurobiological impairments: alterations in functional activity, glucose metabolism, perfusion, and gray matter (GM) atrophy (Lau et al., 2016; Schroeter et al., 2009). GM atrophy is a normal aspect of the aging process (Marie et al., 2023). Results from independent MCI studies and meta-analyses converge towards evident inferotemporal shrinkage, including the amygdala and the hippocampus (Barnes et al., 2009; Ferreira et al., 2011; Nickl-Jockschat et al., 2012; Rao et al., 2023; Schroeter et al., 2009; Tabatabaei-Jafari et al., 2015; Yang et al., 2012; Zhang et al., 2021). The average hippocampal volume decreases linearly from healthy controls (HC) to patients with MCI and dementia (Miao et al., 2022; Schuff et al., 2009). The atrophy of the hippocampus in MCI correlates with cerebrospinal fluid biomarkers (Aβ_1–42_ concentration, (Schuff et al., 2009)), the degree of cognitive impairment (Nickl-Jockschat et al., 2012; Schuff et al., 2009; Xie et al., 2015)) and predicts conversion to dementia (Apostolova et al., 2006; Delli Pizzi et al., 2019; Ferreira et al., 2011; McEvoy et al., 2009; Miao et al., 2022; Rao et al., 2023; Schroeter et al., 2009; Schuff et al., 2009).

Beyond neuroanatomical evaluations, it remains unclear whether hippocampal atrophy in MCI translates into large-scale functional network alterations. Reviews indicate abnormal connectivity in the default-mode network (including the hippocampus), the salience network, the sensorimotor network, the dorsal-attentional network, the executive control and frontoparietal networks, reflecting the various domains affected by MCI (Eyler et al., 2019; Farràs-Permanyer et al., 2015; Kazemi-Harikandei et al., 2022; Song et al., 2021; Xu et al., 2020; Yang et al., 2023). In addition to consistent hypoconnectivity patterns, including, for instance, clusters in the hippocampus and the precuneus (Eyler et al., 2019; Wang et al., 2013), clusters of hyperconnectivity occur and are interpreted as compensatory mechanisms.

Despite numerous reports of widespread functional connectivity alterations in MCI affecting the hippocampus among other brain regions, the association between such alterations and the specific pattern of hippocampal atrophy remains unclear. To the best of our knowledge, only three studies have explored the relationship between hippocampal atrophy and whole-brain functional connectivity using a seed-based approach (Delli Pizzi et al., 2019; Schnellbächer et al., 2020; Xie et al., 2015). Two of them focused on MCI and assessed the relationships among hippocampal atrophy, associated functional connectivity, and behavior (Delli Pizzi et al., 2019; Xie et al., 2015). Yet, such an investigation may provide new insights into the role of the hippocampus in MCI, with implications for diagnosis and potentially, prognosis.

The objective of the present study is to characterize the neuropsychological and sensorimotor profile of MCI and its associated structural and functional brain alterations by combining comprehensive psychometric and sensorimotor testing with voxel-based morphometry (VBM) and seed-based resting-state functional connectivity based on atrophy regions. We examined whether hippocampal atrophy, identified through VBM, would be associated with decreased connectivity of hippocampal networks, and whether both structural and functional indices would relate to the multidomain behavioral impairments observed in this population.

Twenty-five MCI patients were compared with fifteen HC matched for age, education level, and gender. We used linear mixed models to analyze multimodal behavioral scores potentially reflecting decline in MCI. We evaluated whole-brain gray matter atrophy (VBM) in relation with seed-based functional connectivity targeting atrophy regions. We assessed the relationships between brain and behavioral features with pairwise correlation matrices and network-based analysis, with the assumption that those relationships may be impaired in MCI. We hypothesized that MCI patients would perform worse than HC on most cognitive measures, and that significant hippocampal atrophy in MCI would be associated with hypoconnectivity patterns from the hippocampus to frontotemporal and basal ganglia regions. Both structural and functional connectivity indices were expected to show negative correlations with behavior. We expected these three levels of analysis to converge on a common profile in which hippocampal atrophy, functional disconnection of hippocampal networks, and multidomain cognitive and sensorimotor impairment jointly reflect an accelerated and impaired brain-cognition aging trajectory in MCI.

## Materials and Methods

### Randomized controlled trial

This cross-sectional study on the brain and behavioral signature of MCI si exploiting the baseline data of a parent randomized controlled trial, investigating the effects of music and psychomotor interventions in individuals with MCI (baseline data; ClinicalTrials.gov <u>NCT04546451</u>), described elsewhere (James et al., 2023). The full protocol was approved by the Commission cantonale d’éthique de la recherche de Genève (CCER) and the Commission cantonale d’éthique de la recherche sur l’être humain Vaud (CER-VD) and was conducted in accordance with the Declaration of Helsinki. All participants provided written informed consent.

Recruitment for this 2-site study took place at the Memory Center of the Hôpitaux universitaires de Genève (MC-HUG) in Geneva and at the Leenaards Memory Center of the Centre Hospitalier Universitaire Vaudois (LMC-CHUV) in Lausanne, both university-affiliated memory centers. Clinical specialists at the memory centers selected 55- to 80-year-old patients with a clear MCI diagnosis with an MMSE score ≥ 23 (Folstein et al., 1975) or a MoCA score ≥ 17 (Roalf et al., 2013; Trzepacz et al., 2015). Dementia had been formally excluded at recruitment by clinical specialists at both memory centers, based on the integration of cognitive performance and preserved functional autonomy in daily life. Exclusion criteria comprised neurological comorbidities other than MCI (e.g., stroke, epilepsy, established neurodegenerative diagnoses), mental health comorbidities (Hospital Anxiety and Depression Scale score (HADS) > 14, i.e., > 7/21 for anxiety and > 7/21 for depression; (Zigmond and Snaith, 1983)), serious physical and motor comorbidities, impaired or uncorrected hearing impairment, left-handedness, and limited French proficiency. Beyond the initial global cognitive evaluation performed at the clinics, we also administered the Cognitive Telephone Screening Instrument (COGTEL), a multidomain cognitive test, in a face-to-face fashion as a main interest but also a control for dementia (Ihle et al., 2017; Kliegel et al., 2007). None of our MCI patients scored below the dementia cut-off COGTEL of 9.75 (Alexopoulos et al., 2021) (Table 1). Healthy controls met the same inclusion and exclusion criteria except for the MCI diagnosis, they were matched to patients for age, sex, and education level, and were recruited from community-dwelling older adults through local advertisements.

**Table 1.** Demographics of HC and MCI patients at recruitment. No significant differences were found between the HC and MCI groups for age, gender, or education level.

| Matching factors and subtypes | Mean ± standard deviation/number (percentage) | | Group comparison | |
| --- | --- | --- | --- | --- |
|  | HC (N= 15) | MCI (N = 25) | statistic | p-value |
| Age | 70.3 ± 4.6 | 71.5 ± 7.5 | $W = 158.5$ | 0.32 |
| Gender | 13 ♀ (87%) | 18 ♀ (69%) | $\chi^2 = 1.6$ | 0.21 |
| Education level | 3.7 ± 1.3 | 3.2 ± 1.4 | $\chi^2 = 2.9$ | 0.58 |
| COGTEL Weighted Score | 31.9 ± 7.8 | 22.0 ± 7.5 | $t = 4.0$ | $p < 0.001$ |
| Amnesic subtype |  | 8 (32%) |  |  |
| Non-amnesic subtype |  | 15 (60%) |  |  |
| Subjective cognitive decline |  | 2 (8%) |  |  |

### Participants

From the 48 participants initially recruited (32 MCI patients and 16 healthy controls), one MCI patient and one control were excluded after the MRI session due to incidental neurological findings, and five patients withdrew before data collection for personal reasons. The final sample consisted of 15 healthy controls and 26 patients. Subtype information was available for all patients. Across sites, eight individuals (32%) were classified as amnestic MCI (including two with mixed amnestic-dysexecutive or multidomain profiles) and 15 (60%) as non-amnestic MCI (including dysexecutive, attentional, and multidomain presentations). Two individuals (8%) were referred with subjective cognitive concerns without clear objective impairment but were retained in the cohort because they were recruited through the same clinical pathway and met all study inclusion criteria. Complete behavioral data were obtained from all healthy controls and 25 of the 26 patients. Structural MRI data included 25 patients (one excluded for excessive motion), and functional MRI analyses included 24 patients (two excluded for excessive motion).

### Behavioral data collection

#### Cognitive-sensorimotor test battery and behavioral variables used for statistical analyses

Standardized tests were chosen based on their sensitivity to MCI symptoms (James et al., 2023). The order of the tests was pseudorandomized across individuals and timepoints. The initially included clock drawing test (Aprahamian et al., 2009) was excluded from the analyses due to potential test-retest effects related to its inclusion in the MoCA for MCI diagnosis at the CHUV.

#### COGTEL multidomain cognitive test (COGTEL weighted score, COGTEL subtest scores)

The COGTEL (Ihle et al., 2017; Kliegel et al., 2007) evaluates verbal short- and long-term memory (pairs of words) (VSTM, VLTM), working memory (backward digit span, BDS), verbal (letter and categorical) fluency (CF, LF), inductive reasoning, and prospective memory, yielding subtest scores and a weighted score.

#### Binaural speech-in-noise perception (-binaural SRT)

The International Matrix Test (Kollmeier et al., 2015) assesses speech-in-noise (SIN) perception (Worschech et al., 2022; Worschech et al., 2021). The binaural Speech Reception Threshold (SRT, dB) was extracted and inverted (*-1) to align higher scores with better performance. Speech perception intelligibility (speech perception accuracy without noise) was measured to exclude participants with poor hearing (intelligibility <80%) from the speech-in-noise analyses (Metselaar et al., 2008).

#### Processing speed and attention (-TMT A Time, -TMT B Time)

The Trail Making Test parts A and B evaluate processing speed and cognitive flexibility, respectively (Strauss et al., 2006). Times were inverted (*-1) to standardize interpretation across tests, such that higher scores indicated better performance (-TMT A Time, -TMT B Time).

#### Attention (D2 CI)

The D2-R evaluates selective and sustained attention (Brickenkamp et al., 2010, 2015). We analyzed the Concentration capacity Index (CI = number of perceived targets - (number of omission errors + confusion errors)).

#### Manual dexterity and coordination (PP Total Score (TS), Left/Right/Both Hands, Assembly Subtest Scores)

The Purdue Pegboard (Lafayette, 1999) assesses visuo-manual aptitude. We extracted left hand, right hand, both hands, and assembly subtest scores, and analyzed the total score (sum, PP TS).

#### Unipedal balance (UBT, left UBT time, right UBT Time)

The *Unipedal Balance Test (UBT)* evaluates postural balance and leg strength (Bohannon and Tudini, 2018). Participants stand on each foot separately as long as possible, twice in succession on each leg (best time retained on two attempts, left and right foot UBT time).

#### Motor Inhibition (GoNoGo d’)

The GoNoGo assesses motor inhibition (Enge et al., 2014). Sensitivity (d-prime) was computed as the difference between Z-transformed hit and false alarm rates.

### MRI data acquisition and preprocessing

MRI data were acquired on 3.0 T Siemens MRI Prisma fit scanners (Siemens, Erlangen, Germany) at both sites: the Brain and Behaviour Laboratory (Centre Médical Universitaire (CMU) and the Laboratoire de Recherche en Neuroimagerie, CHUV (Centre Hospitalier Universitaire Vaudois, University Hospital of Lausanne).

#### Structural imaging and voxel-based morphometry preprocessing

T1-weighted structural images were acquired using an MP2RAGE sequence (duration = 8:22 minutes; voxel size = 1 mm^3^ isotropic; 176 slices; FOV = 256×240×176 mm; TR/TE = 5000/2.98 ms; TI1/TI2 = 700/2500 ms; flip angle 1/2 = 4°/5°). Preprocessing and statistical analyses were performed using MATLAB R2023a (v9.14.0.2206163), the statistical parametric mapping software (SPM12 r7771, Statistical Parametric Mapping, Wellcome Department of Imaging Neuroscience, London, UK, https://www.fil.ion.ucl.ac.uk/spm/, (Penny et al., 2011) and the Computational Anatomy Toolbox (CAT12, version 12.8.2, r2170, https://neuro-jena.github.io/cat/). Gray Matter (GM) volume maps were extracted using the CAT12 cross-sectional advanced voxel-based morphometry (VBM) pipeline (Ashburner and Friston, 2000; Gaser et al., 2022; Gaser and Kurth, 2017; Marie and Golestani, 2016). Standard CAT12 preprocessing included denoising, tissue segmentation into GM, WM, and CSF, and computation of Total Intracranial Volume (TIV). Segmented images were normalized to MNI space using DARTEL (Ashburner, 2007), modulated based on Jacobian determinants, and smoothed with an 8-mm FWHM Gaussian kernel. Images were statistically checked for homogeneity and quality (CAT12 retrospective quality assurance framework).

#### Resting-state functional MRI preprocessing for seed-based functional connectivity

Resting-state fMRI data were acquired using an EPI sequence (duration: 10:31 min; voxel size = 2.5 mm isotropic; FoV = 210×210×135 mm³; multiband factor = 3; TR = 1350 ms; TE = 31.6 ms). Participants kept their eyes open and focused on a fixation cross. Preprocessing and analysis were performed using the CONN toolbox (RRID:SCR_009550, release 22.a; https://web.conn-toolbox.org; (Nieto-Castanon, 2020)) on SPM12 and Matlab R2023a. The pipeline included realignment with susceptibility distortion correction, slice-timing correction, outlier detection, segmentation, and MNI normalization. Images were smoothed with a 5-mm FWHM Gaussian kernel. Motion artefacts were removed using the Friston 24-parameter model (Friston et al., 1996). WM, CSF, and global signal were regressed out. Data were bandpass filtered (0.008-0.09 Hz).

### Statistical analysis

Statistics were performed using MATLAB R2023a, SPM12, CAT12, and CONN (versions as stated above), as well as JASP (0.18.2). Age and Site (1: Geneva, 2: Lausanne) were added as covariates in all analyses. Analyses involving gray matter volume were additionally controlled for TIV.

#### Demographics and matching

Descriptive statistics (Table 1) report the matching process and dropouts. Continuous variables (Age, COGTEL Weighted Score) were compared using Student t-tests or Mann-Whitney U tests after normality evaluation. Categorical variables were assessed with χ2 tests (education level, gender).

### Behavioral, brain structural and functional signature of MCI compared to healthy aging

#### Behavioral analyses

Binaural SRT, TMT A Time, and TMT B Time were multiplied by −1 so that higher scores consistently reflected better performance. All scores were z-transformed (mean = 0, SD = 1) before modeling, allowing direct comparison between scores of different magnitudes (accuracy, time etc.).

To evaluate cognitive differences between MCI patients and healthy controls across various domains, we fitted a linear mixed-effects model (346 observations, 40 participants) with z-transformed performance as dependent variable and Subject coded as random effect. Fixed effects included Group (MCI vs. HC), Test (9 levels), and their interaction (Group * Test). Between-group differences are reported as Hedges’ g, a standardized effect size measure that corrects for small-sample bias (Cumming, 2013; Hedges and Olkin, 2014).

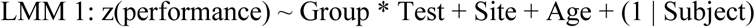

To further investigate the contribution of individual COGTEL subtests, we used the same LMM approach but replaced the Test variable with COGTEL Subtest (6 levels). This analysis included 240 observations from 40 participants.

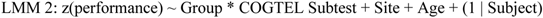

The Prospective Memory subtest, which uses binary scoring (0 = fail, 1 = success), was assessed separately using a χ² test.

#### VBM analyses

To evaluate group differences in GM volume, we performed a whole-brain two-sample *t*-test in SPM (HC vs. MCI) based on smoothed and modulated images. We present results using a statistical threshold of p < 0.001 uncorrected (unc.) at peak-level and p < 0.05 family-wise error corrected (FWEcorr) at cluster-level (k = 150 voxels). Uncorrected results are presented to provide a broader perspective. Anatomical labeling was performed based on neuroanatomical knowledge and the AAL atlas (Tzourio-Mazoyer et al., 2002).

#### Seed-based functional connectivity analyses (atrophy-based functional connectivity)

Functional connectivity strength was represented by Fisher-transformed bivariate correlation coefficients from a weighted general linear model (Nieto-Castanon, 2020). We selected three seeds based on the VBM atrophy clusters, an approach previously applied in MCI and other neurological conditions (Cruz Gómez et al., 2013; Jia et al., 2024; Xie et al., 2015). Those seeds corresponded to the three clusters showing significant atrophy at p < 0.05 FWE corrected at cluster level: the left hippocampus (k = 661), right hippocampus (k = 1029), and anteromedial cerebellum (k = 607 voxels). Group differences were examined using HC > MCI contrasts. We report results at p < 0.005 uncorrected at peak-level and p < 0.05 false discovery rate (FDR) corrected at cluster-level.

#### Relationships between brain, behavior features and age in MCI and HC

To further chart the neuropsychological signature of MCI, we evaluated relationships between hippocampal structural and functional measures, behavioral performance, and Age using two complementary approaches: pairwise partial correlations in the full sample with Age partialled out, and network-based analyses of conditional dependencies among all features including Age, separately in each group.

#### Correlation matrix analysis

We computed pairwise non-parametric partial correlations (Spearman’s rho) between left hippocampal GM volume (L Hip GM), mean functional connectivity of left (L Net FC) and right (R Net FC) networks associated with the left hippocampus, and performance on nine cognitive and sensorimotor tests (Binaural SRT, TMT A, TMT B, COGTEL WS, D2 CI, PP TS, Left UBT, Right UBT, and GoNoGod’), with Age partialled out. GM volume was extracted using the SPM get_totals function applied to individual modulated GM maps and a binary mask of the hippocampal atrophy cluster. Mean functional connectivity values were extracted from CONN.

#### Network-based analysis

Pairwise associations may reflect shared variance rather than independent relationships and differ between MCI and HC. We therefore conducted network analyses estimating conditional dependencies among brain and behavioral features and Age, separately in each group. Edges were estimated using nonparanormal partial correlations while accommodating non-Gaussian distributions (semiparametric Gaussian copula model). Age was included to evaluate whether it was associated with MCI symptoms. We applied a correlational threshold of 0.2 to eliminate very weak associations. The network consisted of 13 nodes (abbreviations as defined in the Behavioral data collection and Correlation matrix analysis sections): (1) GoNoGo d’, (2) Right UBT, (3) Left UBT, (4) PP TS, (5) Binaural SRT, (6) D2 CI, (7) TMT A, (8) TMT B, (9) COGTEL WS, (10) R Net FC, (11) L Net FC, (12) L Hip GM, and (13) Age.

The first analysis yielded zero Closeness values in the MCI network, potentially because GoNoGo d’, the motor inhibition measure, formed an isolated node disconnected from the rest of the network.To further test this assumption, we conducted a second network-based analysis excluding this variable.

Finally, to further characterize relationships between COGTEL subtests and hippocampal features, we conducted a third network-based analysis comprising: (1) CF, (2) LF, (3) BDS, (4) VSTM, (5) VLTM, (6) R Net FC, (7) L Net FC, (8) L Hip GM, and (9) Age.

## Results

### Population demographics and matching factors between MCI and HC

#### Behavioral signature of MCI patients compared to HC participants

The LMM evaluating differences between MCI and HC in various cognitive and sensorimotor domains showed a significant main effect of Group [F(1, 36.13) = 28.8, p < 0.0001], indicating a robust association between group level and performances across all nine tests (-Binaural SRT, -TMT A, -TMT B, COGTEL WS, D2 CI, PP TS, Left UBT, Right UBT, GoNoGo d’). Site and Group * Test interaction were not significant (Figure 1A). Contrasts (HC - MCI, Table 2) were computed for each test based on z-score estimated marginal means (p-values adjusted with the Holm correction), revealing robust impairments in MCI (−0.83 to −1.21 SD from the population mean, Table 2). Age had a significant negative effect on performance [F(1, 36.47) = 6.0, p < 0.02], confirmed by a negative correlation (Spearman’s rho = −0.32, p < 0.0001).

**Figure 1.**
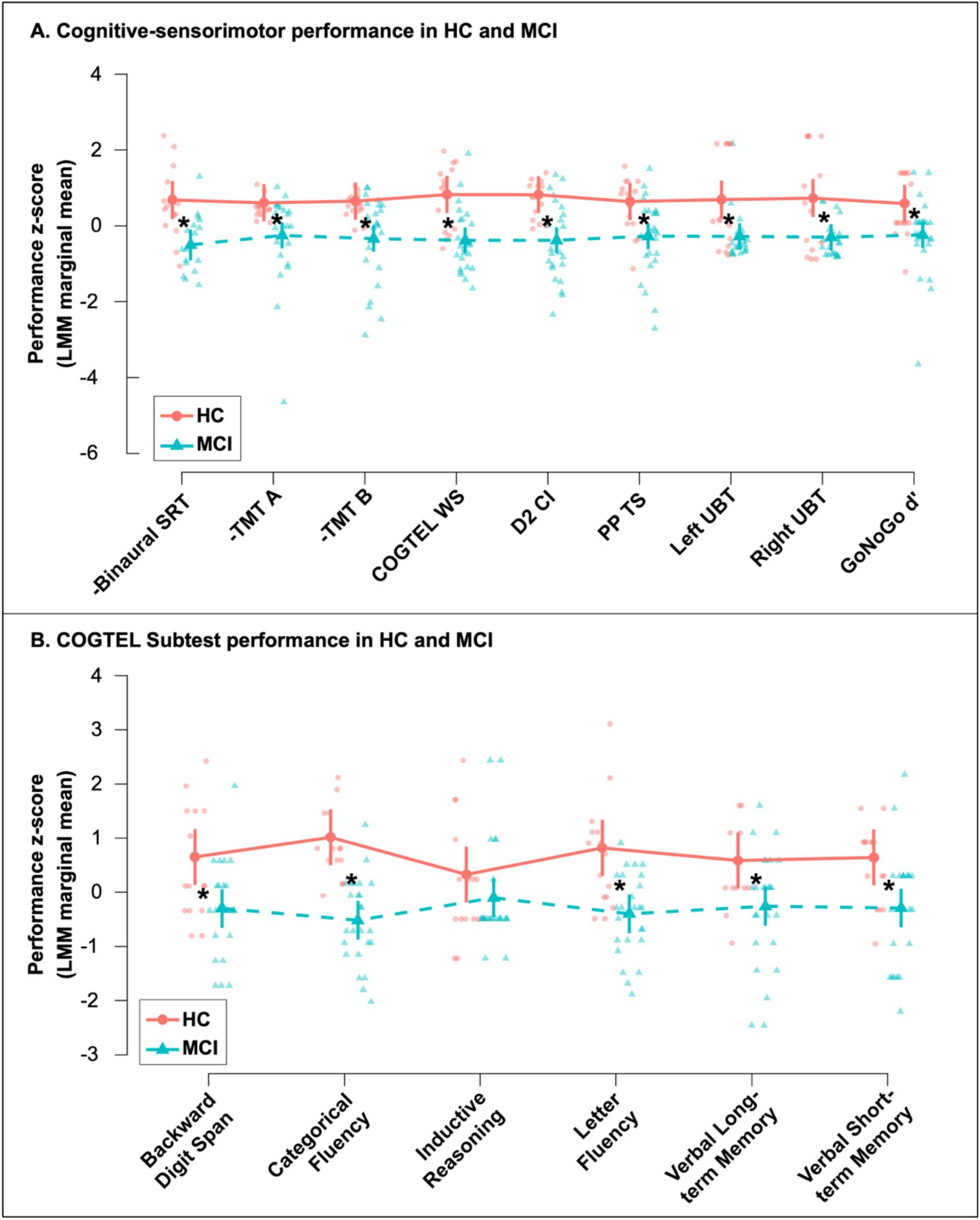
A. Estimated marginal mean z-scores for each cognitive-sensorimotor test in HC and MCI. B. Estimated marginal mean z-scores for each COGTEL subtest in HC and MCI (HC: red, MCI: cyan; asterisks indicate significant group differences).

**Table 2.**
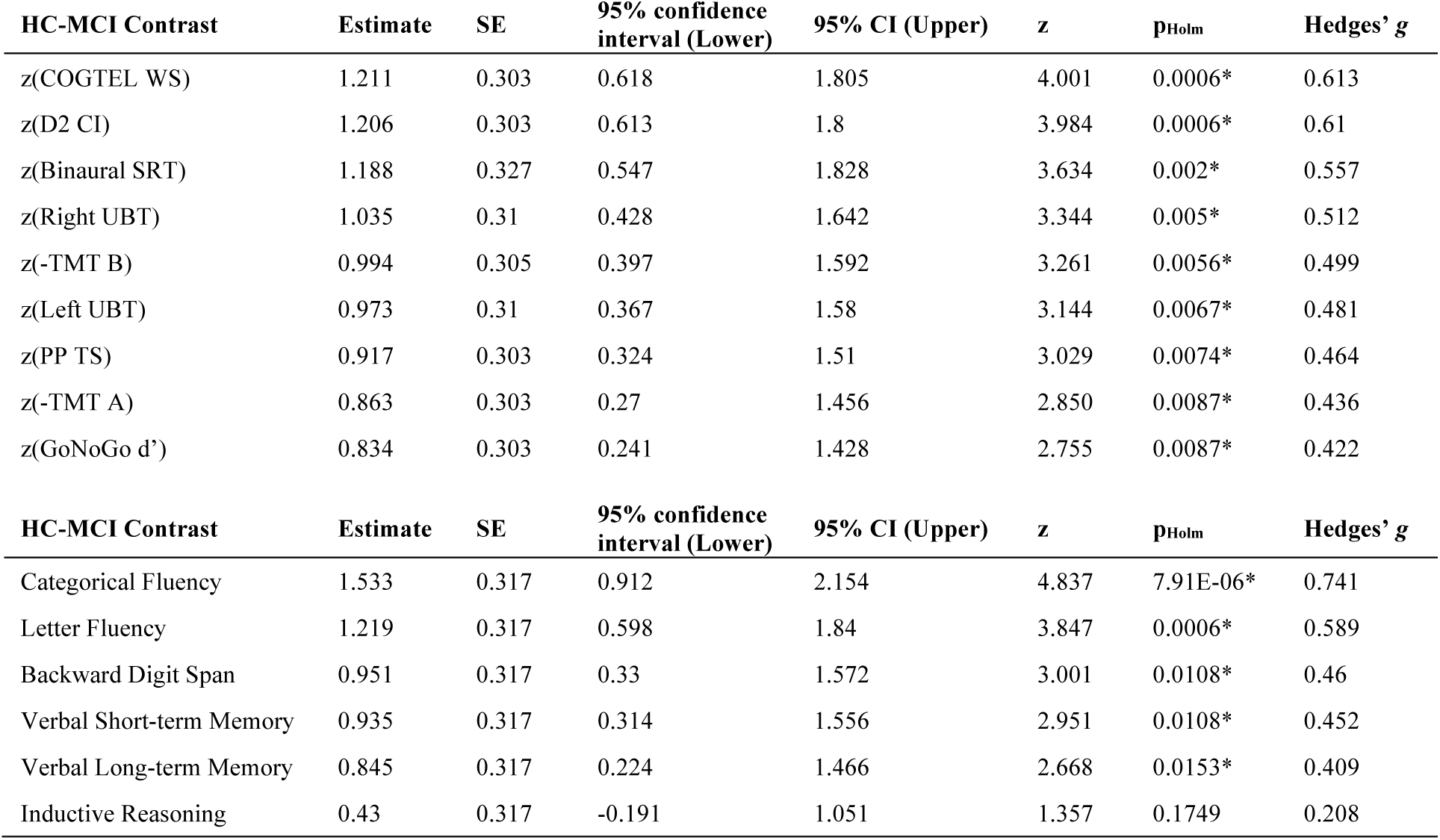
HC vs. MCI estimated marginal means contrasts for each psychometric test and COGTEL subtest performance z-score (2 separate linear mixed models). Effects are presented in descending order of Hedges’ g. P-values are Holm-corrected. SE: standard error; SRT: Speech Reception Threshold; TMT: Trail Making Test; WS: weighted score; CI: Concentration Index; PP TS: Purdue Pegboard Total Score; UBT: Unipedal Balance Test.

| HC-MCI Contrast | Estimate | SE | 95% confidence interval (Lower) | 95% CI (Upper) | z | p<sub>Holm</sub> | Hedges' g |
| --- | --- | --- | --- | --- | --- | --- | --- |
| z(COGTEL WS) | 1.211 | 0.303 | 0.618 | 1.805 | 4.001 | 0.0006* | 0.613 |
| z(D2 CI) | 1.206 | 0.303 | 0.613 | 1.8 | 3.984 | 0.0006* | 0.61 |
| z(Binaural SRT) | 1.188 | 0.327 | 0.547 | 1.828 | 3.634 | 0.002* | 0.557 |
| z(Right UBT) | 1.035 | 0.31 | 0.428 | 1.642 | 3.344 | 0.005* | 0.512 |
| z(-TMT B) | 0.994 | 0.305 | 0.397 | 1.592 | 3.261 | 0.0056* | 0.499 |
| z(Left UBT) | 0.973 | 0.31 | 0.367 | 1.58 | 3.144 | 0.0067* | 0.481 |
| z(PP TS) | 0.917 | 0.303 | 0.324 | 1.51 | 3.029 | 0.0074* | 0.464 |
| z(-TMT A) | 0.863 | 0.303 | 0.27 | 1.456 | 2.850 | 0.0087* | 0.436 |
| z(GoNoGo d') | 0.834 | 0.303 | 0.241 | 1.428 | 2.755 | 0.0087* | 0.422 |
| HC-MCI Contrast | Estimate | SE | 95% confidence interval (Lower) | 95% CI (Upper) | z | p<sub>Holm</sub> | Hedges' g |
| Categorical Fluency | 1.533 | 0.317 | 0.912 | 2.154 | 4.837 | 7.91E-06* | 0.741 |
| Letter Fluency | 1.219 | 0.317 | 0.598 | 1.84 | 3.847 | 0.0006* | 0.589 |
| Backward Digit Span | 0.951 | 0.317 | 0.33 | 1.572 | 3.001 | 0.0108* | 0.46 |
| Verbal Short-term Memory | 0.935 | 0.317 | 0.314 | 1.556 | 2.951 | 0.0108* | 0.452 |
| Verbal Long-term Memory | 0.845 | 0.317 | 0.224 | 1.466 | 2.668 | 0.0153* | 0.409 |
| Inductive Reasoning | 0.43 | 0.317 | -0.191 | 1.051 | 1.357 | 0.1749 | 0.208 |

A subsequent LMM on COGTEL subtests showed a significant main effect of Group [F(1, 36.00) = 21.6, p < 0.0001]. Age and Site effects were not significant. Figure 1B illustrates impaired performance in MCI compared to HC across all subtests except Inductive Reasoning, leading to a marginal Group * COGTEL Subtest interaction [F(5, 190.00) = 1.9, p = 0.08]. Contrasts (HC - MCI, Table 2) showed significant impairment in MCI (−0.84 to 1.53 SD) in each COGTEL subtest except for Inductive Reasoning. A significant decrease in the proportion of MCI patients who successfully completed the prospective memory subtest was also observed compared with HC (20% vs. 60%; χ² = 6.6, p = 0.01).

Together, these results confirm that MCI is associated with a broad profile of cognitive and sensorimotor impairment in this sample, with semantic fluency, attention, and speech-in-noise perception showing the largest group differences between HC and MCI.

#### Voxel-based morphometry signature of MCI

GM volume atrophy analyses (HC > MCI contrast) revealed significant reduction in MCI (Table 3, Figure 2, p < 0.001 uncorrected at peak-level, k = 150 voxels) in 15 fronto-temporal regions. The most significant decrease was observed in three clusters (p < 0.05 FWE at cluster level): the left and right hippocampus, and an anteromedial cerebellar region. The reverse contrast (MCI > HC) yielded no significant results. These atrophy clusters served as seeds for the subsequent functional connectivity analyses.

**Figure 2.**
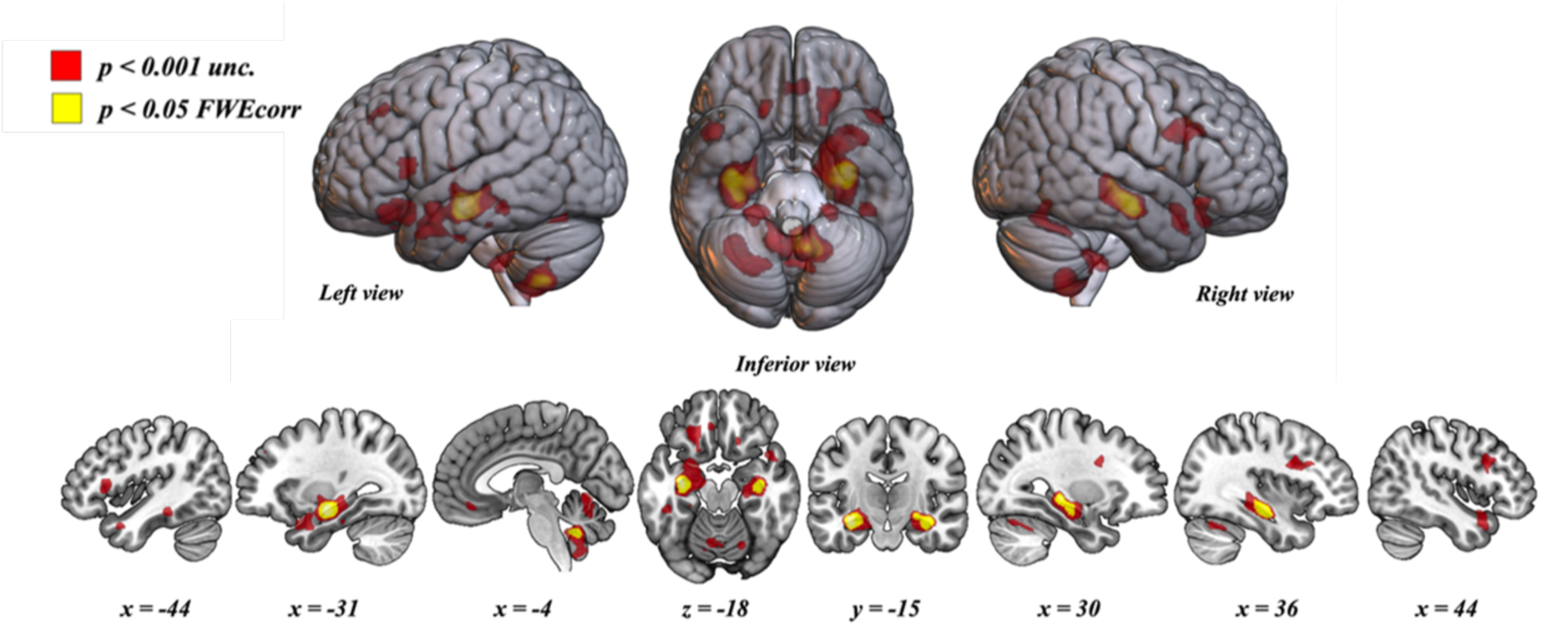
G Gray matter volume decrease in MCI compared to HC (HC > MCI), displayed on the MNI152 brain template (red: p < 0.001 uncorrected at peak-level; yellow: p < 0.05 family-wise error corrected at cluster-level; k = 150 voxels).

**Table 3.**
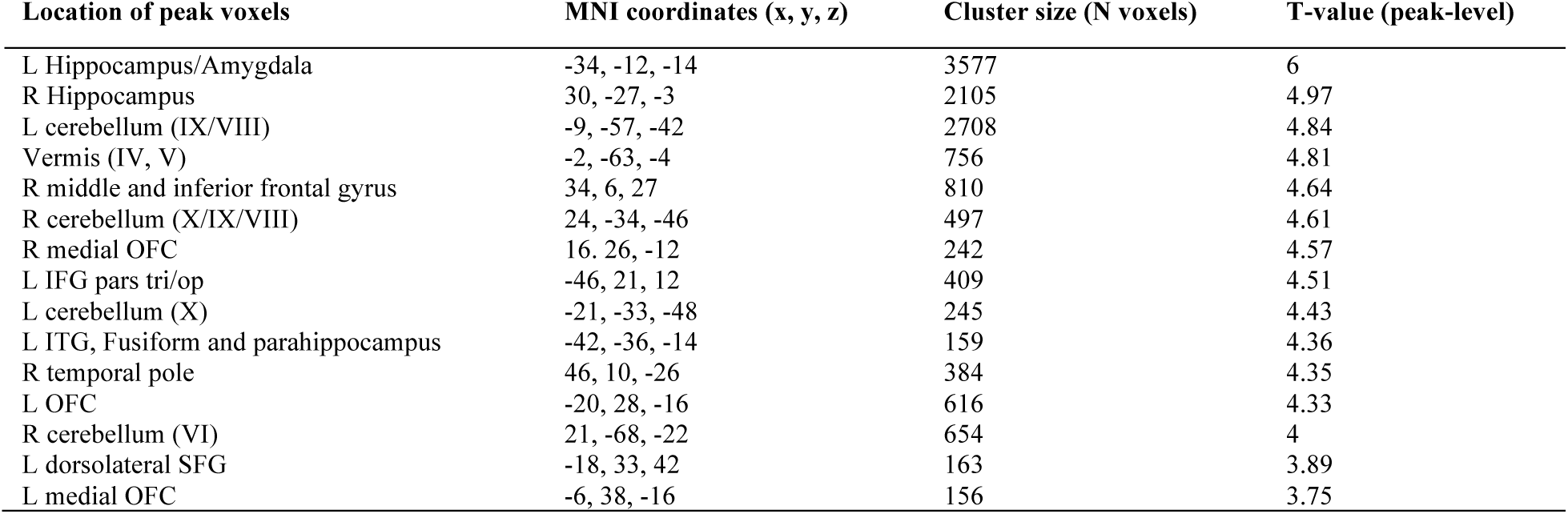
Gray matter volume reduction in MCI compared to HC (p < 0.001 uncorrected at peak-level, k = 150 voxels). The first three clusters are significant at p < 0.05 FWE-corr. at cluster-level. L: left; R: right; OFC: orbitofrontal cortex; tri: triangularis; op: opercularis; SFG: superior frontal gyrus; MNI: Montreal Neurological Institute.

#### Functional connectivity signature of MCI

Within-group analyses revealed significant positive functional connectivity between the left hippocampal atrophy cluster and widespread cortical and subcortical regions (Figure 3A). In HC, functional connectivity extended bilaterally across fronto-temporo-parietal cortices, including premotor and motor areas, the precuneus, insula, and cingulate cortex, and subcortically with the basal ganglia and thalamus. In MCI, a qualitatively similar but spatially reduced pattern was found, with smaller clusters in homologous cortical regions and more restricted subcortical connectivity (Figure 3, Table 4). Within-group analyses were conducted for all three seeds but are shown only for the left hippocampal seed, which was the only seed yielding significant group differences.

**Figure 3.**
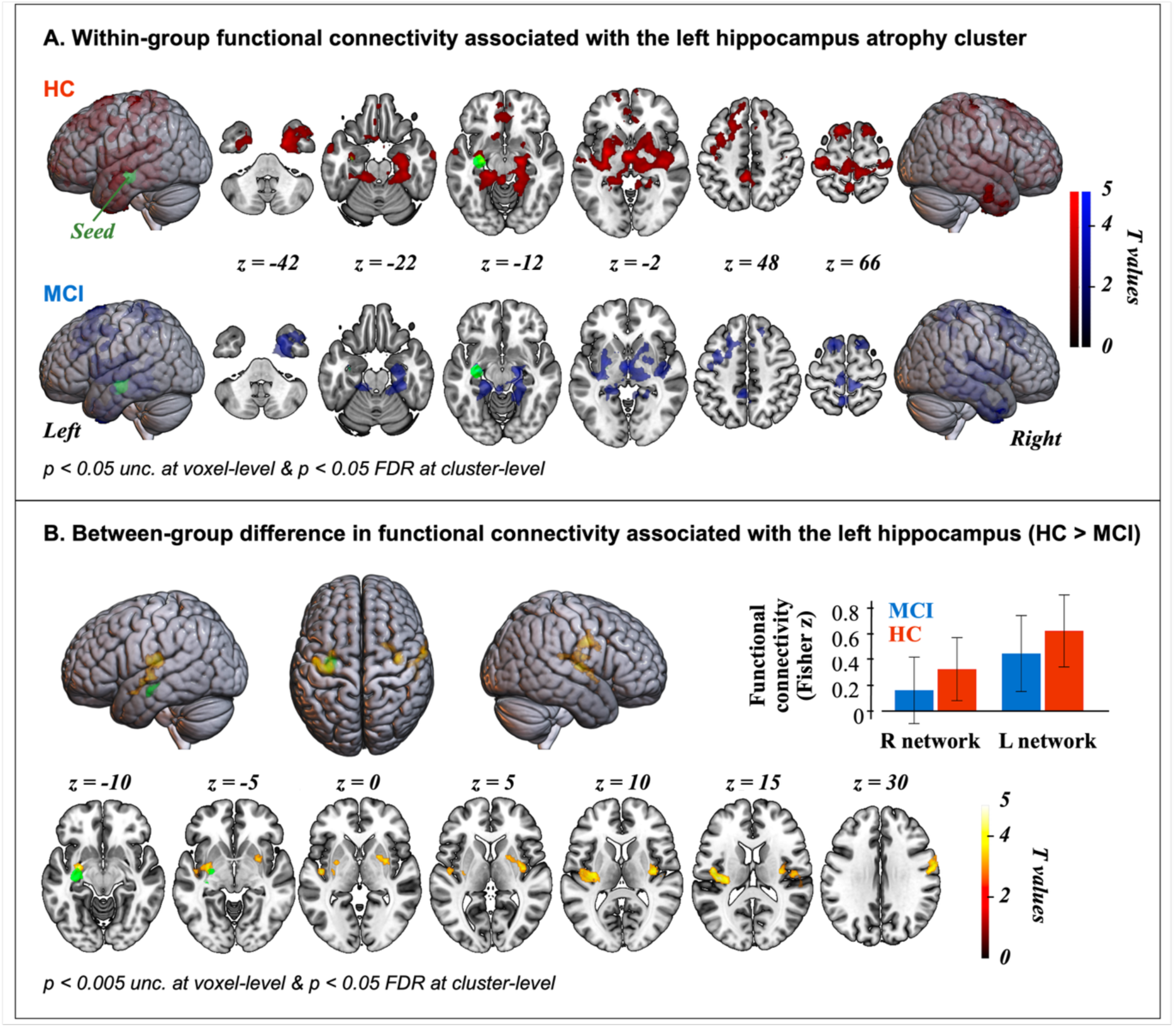
Functional connectivity associated with the left hippocampal atrophy seed (green) in A. HC (red) and MCI (blue), and B. HC > MCI contrast (hot color scale), displayed on the MNI152 brain template.

**Table 4.**
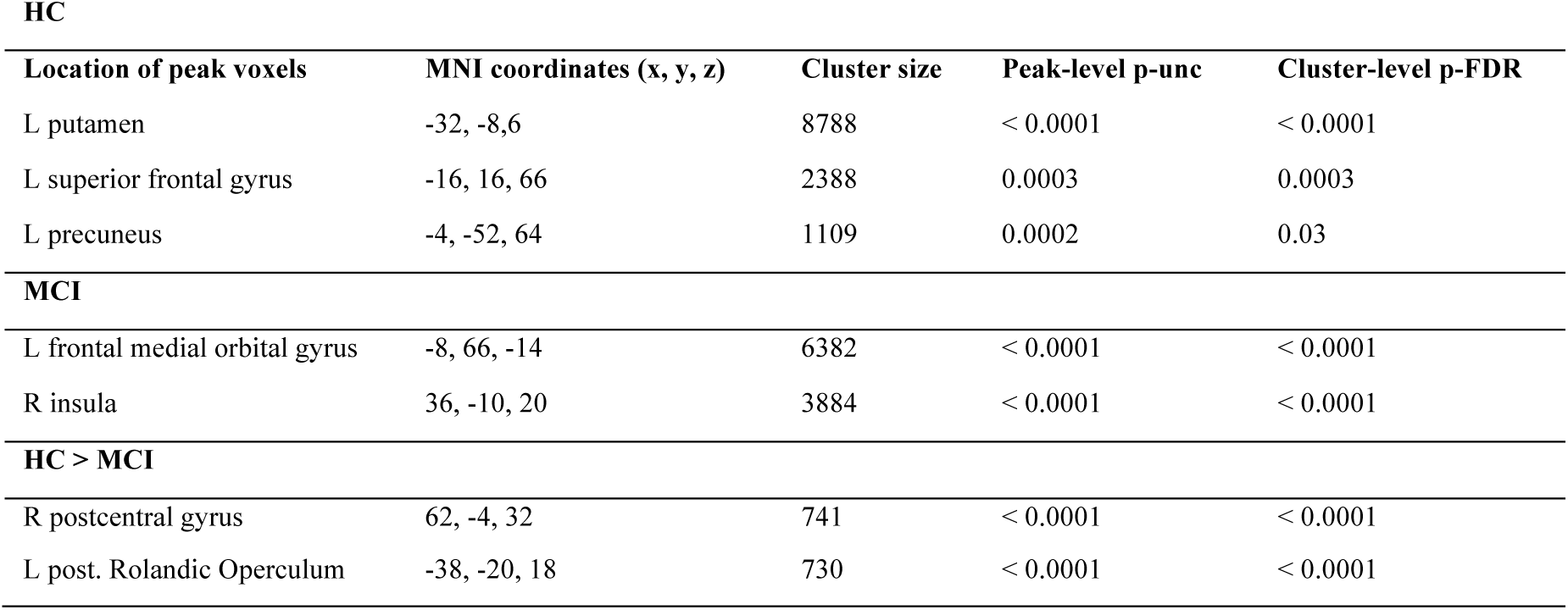
Functional connectivity clusters associated with the left hippocampal atrophy seed in HC, MCI, and HC > MCI contrast. L: left; R: right; post: posterior.

The HC > MCI contrast revealed reduced functional connectivity in MCI in sensorimotor and executive control regions, primarily distributed around the bilateral Sylvian fissure including the insula, Rolandic operculum, putamen, and pallidum, as well as the left orbital frontal cortex and pars orbitalis of the inferior frontal gyrus, and right superior frontal gyrus (Figure 3, Table 4, p < 0.005 uncorrected at peak-level, p < 0.05 false discovery rate corrected at cluster-level). The right hippocampal and anteromedial cerebellar seeds did not yield significant group differences.

Left hippocampal atrophy was thus associated with reduced functional connectivity in bilateral sensorimotor, insular, striatal, and frontal regions, while right hippocampal and cerebellar atrophy yielded no functional connectivity differences.

### Relationships between brain and behavioral features of MCI

#### Independent relationships in the full population of MCI and HC participants

In the full sample (MCI and HC), the age-adjusted partial correlation matrix (Figure 4) shows predominantly positive associations between brain and behavioral measures. Structural and functional hippocampal markers exhibit strong intercorrelations (Spearman’s ρ ≈ 0.65–0.83). Most cognitive and sensorimotor measures correlate positively with these brain metrics (ρ ≈ 0.33–0.83), with category ical fluency showing the highest associations. In contrast, balance and GoNoGo d′ show weak associations with brain variables.

**Figure 4.**
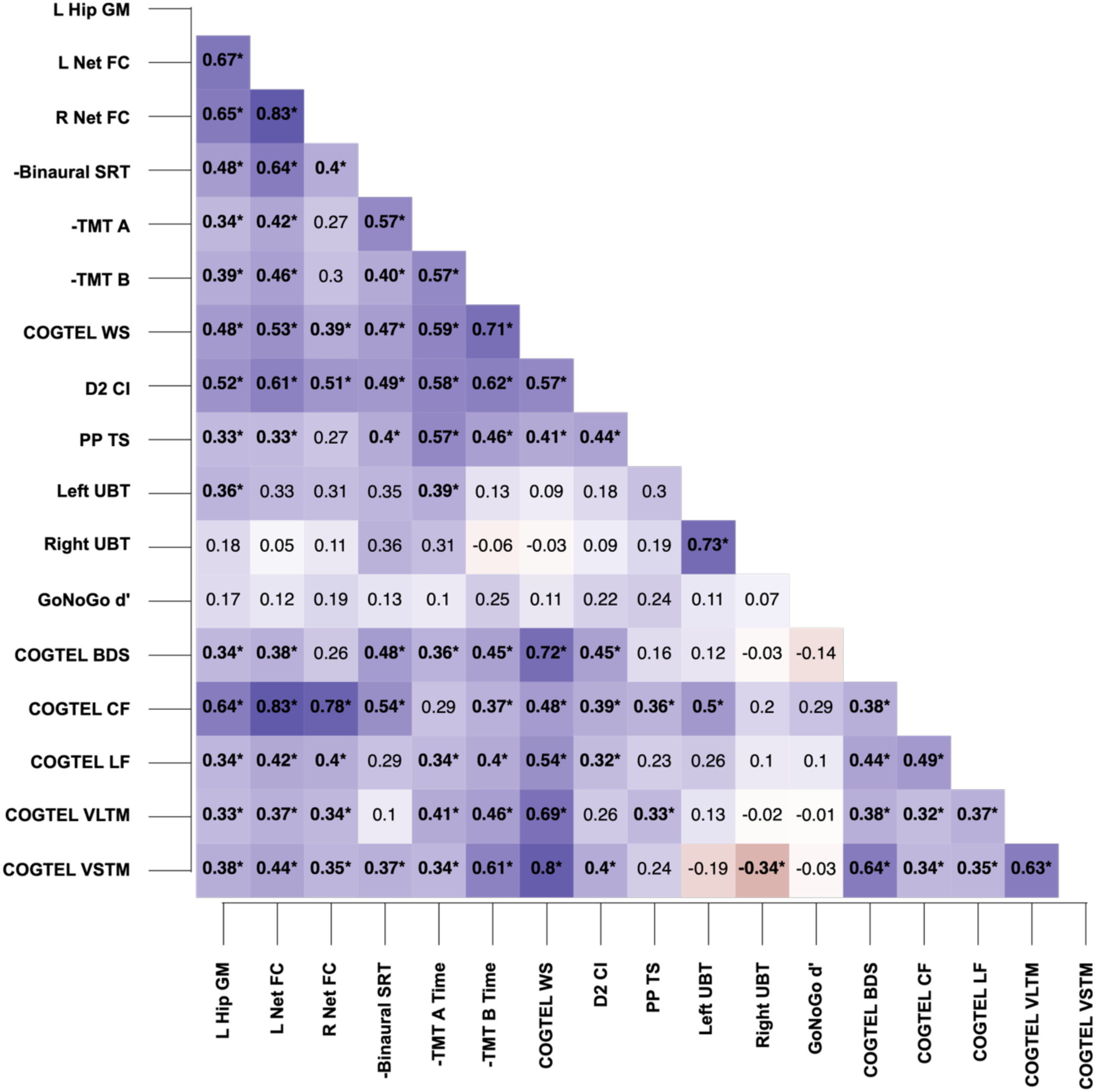
Age-adjusted partial correlation matrix between hippocampal structural and functional measures and cognitive-sensorimotor performance in the full sample (MCI and HC). Significant correlations are indicated by asterisks and bold font. L Hip GM: left hippocampal gray matter volume; L/R Net FC: left/right network functional connectivity associated with left hippocampal atrophy; SRT: Speech Reception Threshold; TMT: Trail Making Test; WS: weighted score; CI: Concentration Index; PP TS: Purdue Pegboard Total Score; UBT: Unipedal Balance Test; BDS: Backward Digit Span; CF: Categorical Fluency; LF: Letter Fluency; VLTM: Verbal Long-Term Memory; VSTM: Verbal Short-Term Memory.

Across the full sample, hippocampal structure and functional connectivity were both positively associated with cognitive performance, particularly semantic fluency, attention, speech-in-noise perception, and general cognition, but not with balance and motor inhibition.

#### Network-based relationships between brain and behavior in MCI and HC

Across all three network models (Fig. 5A-C), HC consistently displayed much denser partial correlation networks than MCI (lower sparsity). In the full multimodal network (Fig. 5A), HC showed numerous conditional associations (sparsity = 0.21; 62/78 edges), with L Net FC, Right UBT, and COGTEL WS emerging as the most influential nodes, and TMT B and binaural SRT also occupying central positions. The MCI network was markedly sparser (sparsity = 0.59; 32/78 edges). Compared to HC, MCI networks consistently exhibited reduced connectivity, with null Closeness values in the first network, GoNoGo d′ being fully disconnected (node 1 in Figure 5A). Sparsity was twofold lower in HC than in MCI in model B, and threefold lower in models A and C.

**Figure 5.**
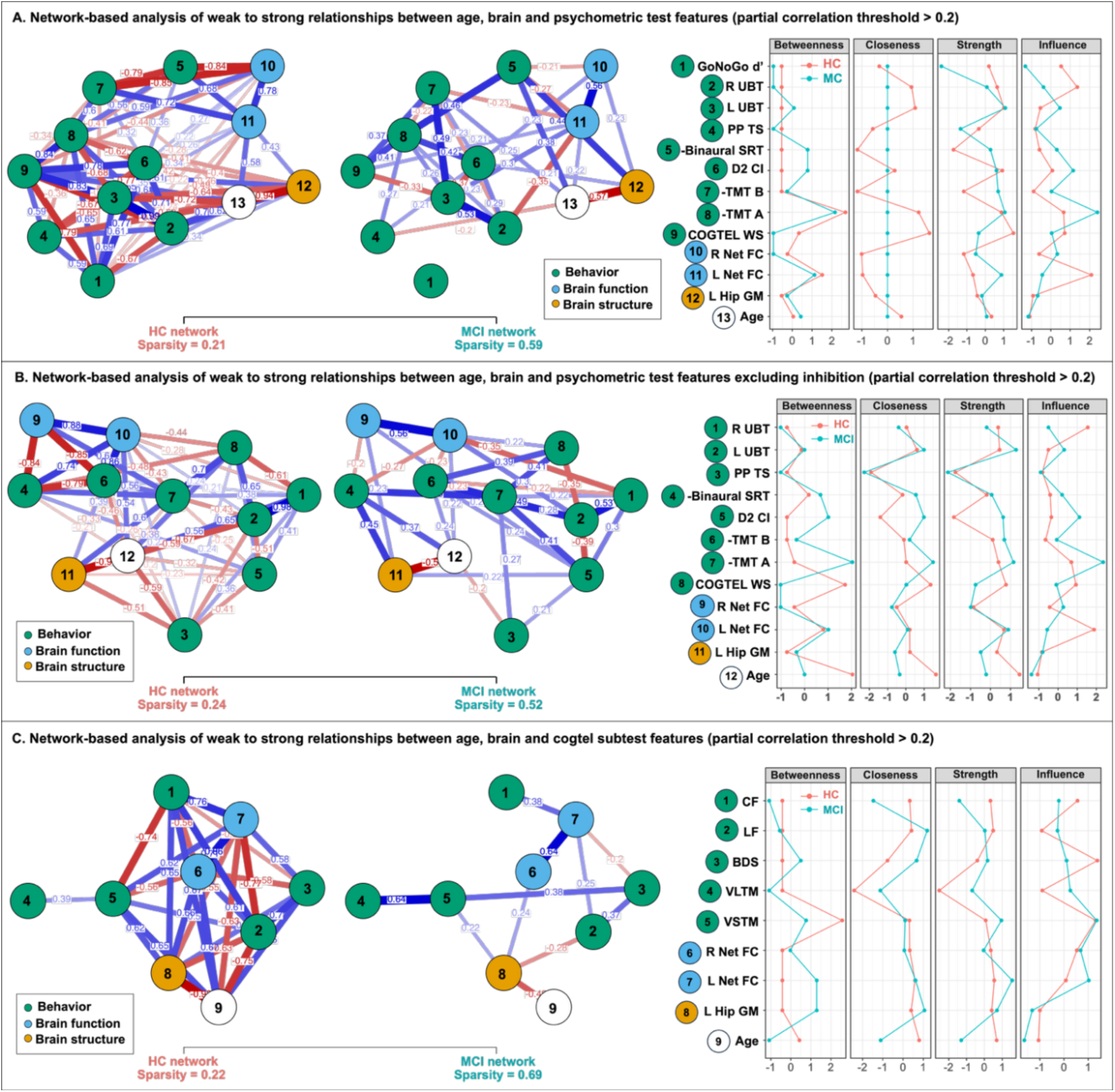
Nonparanormal partial correlation networks of brain, cognitive, and age-related features (correlation threshold = 0.2) in HC (red) and MCI (cyan). A, Full cognitive-sensorimotor, hippocampal structural, and functional feature set. B, Identical model excluding inhibition. C, COGTEL subtests replacing cognitive-sensorimotor tests. Node color indicates feature domain (green = behavior, blue = brain function, orange = brain structure). Edge color reflects correlation sign (blue = positive, red = negative); edge thickness reflects strength. Centrality metrics (betweenness, closeness, strength, expected influence) are plotted for each group. BDS: Backward Digit Span; CF: Categorical Fluency; CI: Concentration Index; L Hip GM: left hippocampal gray matter volume; L Net FC: functional connectivity of the left network associated with left hippocampal atrophy; LF: Letter Fluency; R Net FC: functional connectivity of the right network associated with left hippocampal atrophy; SRT: Speech Reception Threshold; TMT: Trail Making Test; UBT: Unipedal Balance Test; VLTM: Verbal Long-Term Memory; VSTM: Verbal Short-Term Memory; WS: Weighted Score.

The second network excluding GoNoGo d′ (Figure 5B) confirmed that inhibitory control did not account for the group differences: HC networks remained highly connected (sparsity = 0.24; 50/66 edges) with similar node rankings, while the MCI network remained sparse (sparsity = 0.52; 32/66 edges) with reduced global centrality values.

The third network incorporating COGTEL subtests (Figure 5C) yielded nine-node networks. HC showed high density (sparsity = 0.22; 28/36 edges), with BDS and VSTM displaying the strongest influence and closeness. MCI networks remained sparse (sparsity = 0.69; 11/36 edges) and lacked clear hubs. CF showed moderate centrality but substantially lower influence and betweenness than in HC. LF, left hippocampal volume, and L Net FC occasionally approached or exceeded HC values on selected centrality metrics.

Across all models, Age was more central in MCI than in HC, showing higher strength and betweenness, generally through negative associations with brain-behavior features. Taken together, these network models indicate that hippocampal structure and functional connectivity were broadly and densely coupled to cognitive and sensorimotor performance in HC, whereas in MCI this coupling was markedly reduced and largely restricted to verbal fluency, with age emerging as the dominant negative influence on the neuropsychological profile. Conditional dependencies between hippocampal measures and behavioral performance were markedly reduced in MCI compared to HC. Where present, the strongest remaining coupling involved verbal fluency.

## Discussion

The present study set out to characterize the neuropsychological, structural, and functional signature of MCI across amnestic and non-amnestic subtypes, with a novel focus on the multivariate coupling between hippocampal structure, functional connectivity, age, and multidomain cognitive and sensorimotor performance. MCI was associated with widespread cognitive and sensorimotor decline, bilateral hippocampal and cerebellar atrophy, and reduced cortico-subcortical connectivity in networks associated with left hippocampal atrophy. Pairwise analyses revealed positive associations between hippocampal structural and functional measures and multidomain behavioral impairments. Network-based analyses further revealed an accelerated and decoupled brain-cognition aging profile in MCI.

### The neuropsychological profile of MCI

#### Multimodal cognitive and sensorimotor deficits associated with MCI compared to healthy aging

Across domains, MCI impairments ranged from −0.83 to −1.53 SD, a conservative estimate because z-scores were computed on the full population (MCI and HC participants), consistent with literature reporting average impairments of ∼1–2 SD in MCI (Albert et al., 2011; Association, 2013; Bradfield, 2023; Machulda et al., 2019; Petersen et al., 1995). Reported in descending order, the largest effects (g ≈ 0.74–0.5) appeared in categorical and letter fluency, general cognition (COGTEL WS), attention, speech-in-noise perception, right-foot balance, and cognitive flexibility. Moderate impairments (g ≈ 0.49-0.41) were found in left-foot balance, working memory, verbal short-term memory, hand dexterity, bimanual coordination, verbal long-term memory, and inhibition. Finally, inductive reasoning showed a small, non-significant impairment. Overall, the MCI group exhibited a typical pattern of heterogeneous impairment, consistent with reports that multi-domain impairment are common (Nordlund et al., 2005).

#### Alternative cognitive and sensorimotor targets for MCI assessment and training

We investigated a broad range of cognitive domains and, beyond classical deficits, identified less conventional impairments, including binaural speech-in-noise perception (SIN), hand dexterity, bimanual coordination, and balance. Our findings identify verbal fluency, especially semantic fluency, and to a lesser extent phonemic fluency, as the most sensitive cognitive markers of MCI, relative to HC, consistent with prior studies (Bialek and Borkowska, 2024; McDonnell et al., 2020; Nutter-Upham et al., 2008). These impairments often emerge early, making verbal fluency a rapid and efficient screening tool (Cintoli et al., 2024; Gustavson et al., 2020), and a strong candidate for monitoring progression. The brain structural correlates of fluency deficits (Molinuevo et al., 2011) overlap with inferotemporal atrophy characteristic of MCI and Alzheimer’s disease (AD) (Nickl-Jockschat et al., 2012; Yang et al., 2012; Zhang et al., 2021), supporting a neurobiological basis for their diagnostic and predictive value.

Verbal fluency may provide additional diagnostic value. Fluency improves MMSE-based MCI detection (McDonnell et al., 2020) and is among the MoCA’s most discriminative subtests (Cecato et al., 2016). Although MMSE and MoCA offer comparable predictive accuracy (Dubbelman et al., 2024), the occasional advantage of the MoCA (Pinto et al., 2019) may stem from its fluency component. The COGTEL, which correlates strongly with the MMSE (r = .93;(Ihle et al., 2017)), and comprises both phonemic and semantic fluency, showed high sensitivity in our data. The COGTEL multidomain coverage, resistance to ceiling effects, and prognostic value-comparable to the modified MMSE-(Alexopoulos et al., 2020; Alexopoulos et al., 2021; Breitling et al., 2010), make it a valuable complement to standard screening tools. Examining COGTEL subtests individually may further enhance diagnostic precision.

SIN perception, the ability to extract speech under competing noise, is a critical real-life competence. Pure-tone audiometry explains only limited variance in daily hearing (Benzaquén et al., 2025; Demeester et al., 2012; Pronk et al., 2013). SIN declines with age (Pronk et al., 2013) and captures additional auditory top-down cognitive processes, including selective attention, processing speed, executive functions including inhibitory control and working memory (Benzaquén et al., 2025; Dryden et al., 2017; Yumba, 2017). After excluding participants with intelligibility inferior to 80% (Metselaar et al., 2008), we observed expected HC and impaired MCI performance (–5.8 ± 1.4 vs. −4.3 ± 1.1), consistent with SIN deficits in MCI (Lee et al., 2016, 2018; Mamo and Helfer, 2021) and their stronger expression in dysexecutive profiles (Lee et al., 2018).

We report hand-dexterity and balance impairments in MCI, consistent with evidence that sensorimotor impairments accompany or precede cognitive decline (Bahureksa et al., 2016; Delbaere et al., 2012; Lodha and McCarthy, 2021; Tsiakiri et al., 2025). Fine motor dexterity declines progressively from healthy to MCI and AD stages (de Paula et al., 2016).

These markers potentially carry prognostic value. Impaired SIN perception predicts incident dementia (Stevenson et al., 2022) and is associated with adverse outcomes linked to social isolation (Livingston et al., 2024). Balance deficits predict functional decline and increased fall risk (Bahureksa et al., 2016; Wang et al., 2024). Verbal fluency tracks progression from MCI to dementia (Cintoli et al., 2024; Gustavson et al., 2020), Together, these findings support the potential clinical relevance of these measures for identifying and monitoring older adults at risk of cognitive decline.

Our results suggest that combining the COGTEL with measures of attention, flexibility, SIN perception, and whole-body balance may provide an efficient complementary strategy for early MCI detection. This battery requires minimal material, takes about 45 minutes to administer, and showed moderate-to-large group differences in the present study (Hedges’ g ≥ 0.5 for all tasks).

#### Hippocampal and cerebellar atrophy patterns in MCI

Our MCI sample showed predominant gray matter atrophy in the bilateral hippocampi and an anteromedial cerebellar region. The hippocampal finding replicates one of the most consistent hallmarks of MCI and AD (Barnes et al., 2009; Ferreira et al., 2011; Nickl-Jockschat et al., 2012; Rao et al., 2023; Schroeter et al., 2009; Tabatabaei-Jafari et al., 2015; Yang et al., 2012; Zhang et al., 2021). Hippocampal atrophy increases from healthy aging to MCI and AD, relates to CSF biomarkers, and predicts cognitive decline and conversion to dementia (Apostolova et al., 2006; Delli Pizzi et al., 2019; Ferreira et al., 2011; McEvoy et al., 2009; Miao et al., 2022; Rao et al., 2023; Schroeter et al., 2009; Schuff et al., 2009).

Cerebellar atrophy in MCI is a relatively new finding, reported in a few recent studies (Ghiyamihoor et al., 2025; Toniolo et al., 2018; van de Mortel et al., 2021). Van de Mortel et al. described atrophy patterns in a large cohort of 1386 MCI and AD patients, reporting that cerebellar atrophy progresses with the degree of cognitive decline, including early MCI involvement of lobule IX, consistent with the cluster spanning lobules VIII–X observed here. The polymodal neocerebellum (VII–IX) supports associative and executive networks (Habas, 2021), and lobule VIII has been linked to working memory performance in healthy older adults (Marie et al., 2023). Large-scale data from approximately 45,000 participants (Ghiyamihoor et al., 2025) suggest age-dependent cerebellar volume loss, with MCI displaying larger attrition than AD. MRI-based cerebellar features may predict MCI with an accuracy comparable to hippocampal markers (Lu et al., 2025), and cerebellar atrophy may even precede hippocampal changes, suggesting that it could represent an additional early structural marker (van de Mortel et al., 2021).

#### Selective impact of left hippocampal atrophy on functional connectivity in MCI

MCI was associated with gray-matter loss, primarily in both hippocampi and an anteromedial cerebellar region. Only the left hippocampal atrophy was associated with reduced connectivity in bilateral cortico-subcortical networks involving temporal, frontal, opercular, insular, striatal, and thalamic areas, a pattern consistent with the multimodal impairments observed in our population. In contrast, right hippocampal and cerebellar atrophy did not yield detectable functional differences, suggesting structural loss does not systematically translate into hypoconnectivity or that its functional consequences may be buffered by compensatory mechanisms or functional reserve processes. No hyperconnectivity was observed.

These findings converge with those of Schnellbächer et al. (Schnellbächer et al., 2020), who modeled FC networks in a large population from atrophy-based seeds obtained in a previous MCI study (Nickl-Jockschat et al., 2012). Despite our smaller left hippocampal seed, our within-group FC maps closely resemble theirs, with broader frontal-parietal extensions likely due to differing statistical thresholds. Our findings are also consistent with those of Delli Pizzi et al. (Delli Pizzi et al., 2019), who reported hippocampal/entorhinal hypoconnectivity in MCI converters and in patients with AD-related CSF changes. Their hyperconnectivity findings in non-converters did not emerge here, potentially reflecting differences in recruitment (amnestic vs. mixed subtypes) and seed definitions (individual parcellations vs. atrophic regions). Using a methodology closest to ours, Xie et al. (Xie et al., 2015) also identified left hippocampal hypoconnectivity in MCI. However, their broader atrophy map and the presence of both of hyper- and hypoconnectivity differ from our data, potentially due to preprocessing choices (optimized vs. advanced VBM, logit-transformed maps, no TIV or age covariates). The overlap in left hippocampal FC patterns across studies supports the reproducibility of this network alteration in MCI. Further large-scale studies are needed to clarify how local degeneration may be associated with functional disruptions and whether compensatory connectivity emerges at specific disease stages.

#### Impaired relationships between hippocampus structure, functional connectivity, and various cognitive-sensorimotor domains in MCI

Pairwise analyses revealed positive relationships between hippocampal GM volume, associated functional connectivity, and nearly all cognitive and sensorimotor domains. Connectivity correlated more strongly with behavior than gray matter. Functional indices may explain additional cognitive variance in aging and MCI (Franzmeier et al., 2017; Lowe et al., 2019; McIntosh and Lobaugh, 2004). Associations spanned general cognition, memory, executive function, and SIN perception, supporting the involvement of fronto-temporal circuits in verbal fluency deficits in MCI (Vonk et al., 2020; Wright et al., 2023). Network modeling extended these results, providing a multivariate view of conditional dependencies among all variables, consistent with previous evidence linking hippocampal structural and functional alterations to cognitive deficits in amnestic MCI (Delli Pizzi et al., 2019; Xie et al., 2015).

Across networks, MCI showed reduced connectivity and lower centrality among behavioral and neural variables, a loss of dominant hubs, and reduced network integration among memory, executive functions, hippocampal structure, and functional connectivity. These findings align with graph-theoretical and connectomics evidence of disrupted hub organization and efficiency in MCI/AD (Sala-Llonch et al., 2015; Tijms et al., 2013; Yu et al., 2021). Fragmentation persisted after removing inhibition, indicating that reduced conditional associations were not driven by a single domain but reflected a broader reduction in interrelationships across cognitive, sensorimotor, and neural features. The complete disconnection of motor inhibition (GoNoGo d’) from the MCI network may indicate an early and selective decoupling of inhibitory control from hippocampal and cognitive systems, beyond what pairwise analyses reveal. HC networks showed a structured and hierarchical organization, with memory and executive functions, hippocampal measures, and SIN perception emerging as central contributors, in accordance with the critical role of the hippocampus for memory and executive function (Simons and Spiers, 2003), and research on SIN and cognition in aging/MCI (Humes, 2021; Maharani et al., 2018).

Network modeling revealed topological disruptions in MCI that were not visible in pairwise associations, capturing shared variance across features and highlighting the added value of multivariate approaches. Whereas healthy aging exhibited an integrated profile, MCI displayed high sparsity between neural and cognitive features, as well as a stronger adverse age effect. This pattern suggests an accelerated brain-cognition aging in MCI, in accordance with independent findings of stronger age-related constraints on cognition (Albert et al., 2011; Association, 2013; Bradfield, 2023; Machulda et al., 2019; Petersen et al., 1995), brain connectomics (Filippi et al., 2023), and an older brain structural phenotype relative to chronological age (Gaser et al., 2013; Huang et al., 2021; Li et al., 2025). These findings raise the possibility of developping of multimodal aging indices that integrate the association between neural and behavioral features, thereby extending current BrainAge frameworks toward more comprehensive estimators. Such “brain–cognitive age” estimators may bring new perspectives on brain and cognitive reserve and could prove useful for characterizing individual differences in aging and disease progression.

### Towards an integrated structure-function-behavior framework of accelerated brain-cognition aging in MCI

In healthy aging, hippocampal gray matter volume, cortico-subcortical functional connectivity, and cognitive performance form a densely integrated hierarchical system. In MCI, this integration appears selectively disrupted. Structural loss was prominent but not uniformly associated with functional alterations: only the hippocampal atrophy locus was associated with global hypoconnectivity. Furthermore, functional disconnection did not map uniformly onto behavioral deficits. Rather, the coupling between these levels was fragmented and highly domain-specific, persisting most strongly for verbal fluency and general cognition. Jointly, hippocampal atrophy, functional disconnection, and multidomain cognitive decline converge on a common accelerated and decoupled aging profile, in which the normal hierarchical organization of brain and behavior appears progressively lost. Age, rather than hippocampal structure or connectivity, emerged as the most central feature in MCI. Our network data suggest that MCI may involve more than a simple acceleration of normal aging, but a qualitative reorganization of brain-cognition relationships in which hippocampal structure and functional connectivity show reduced predictive value for cognitive performance, and age becomes the dominant determinant of the neuropsychological profile.

### Limitations

The MCI sample was larger than the HC sample to address the longitudinal aspect of the randomized controlled trial not presented here, with sample sizes determined by the power calculation of the parent RCT (James et al., 2023). This imbalance between groups may have limited effects, as healthy populations show less cognitive variability than MCI patients (Anderson et al., 2016) and linear mixed models cope well with unbalanced group sizes (Schielzeth et al., 2020). Findings should be interpreted cautiously and tested for reproducibility in larger cohorts. The cross-sectional design precludes causal inference and limits conclusions about the temporal relationships between hippocampal atrophy, functional disconnection, and cognitive decline. Recruitment through two University Memory clinics, while targeting early-stage MCI (MMSE ≥ 23), may limit generalizability to more advanced or community-detected cases. However, strict inclusion criteria and clinical diagnosis at certified memory centers ensured diagnostic homogeneity across the sample. Selection bias cannot be excluded either, as less impaired individuals may have chosen to participate.

Although functional connectivity seeds were defined from atrophy clusters identified in the present sample, structural and functional analyses rely on independent imaging modalities, an approach previously validated in MCI and other neurological conditions (Cruz Gómez et al., 2013; Jia et al., 2024; Xie et al., 2015). The absence of functional connectivity differences for two of the three atrophy clusters further supports the notion that structural results do not artificially generate connectivity findings. Finally, amyloid biomarker confirmation was performed systematically at one recruitment site but not at the other, where it was conducted only when clinically indicated, preventing formal biological staging of MCI across the full sample.

## Conclusion

Our findings indicate that MCI is characterized by multidomain cognitive and sensorimotor vulnerabilities, with semantic fluency, executive function, and speech-in-noise perception emerging as sensitive, complementary markers. Hippocampal atrophy and reductions in connectivity, together with behavioral impairments and aging, are consistent with a pattern of accelerated and decoupled brain-cognition aging in MCI. The cognitive-sensorimotor battery used here, including the COGTEL, and multimodal aging indices integrating neural and behavioral features, may complement existing diagnostic tools for MCI.

Several affected domains are modifiable through targeted non-pharmacological interventions. Music practice and psychomotor training are promising candidates, with documented transfer effects to cognition, sensorimotor function, and brain plasticity in older adults and MCI (Dorris et al., 2021; Ito et al., 2022; James et al., 2024; Jordan et al., 2022; Martins et al., 2024; Pereira et al., 2018; Raglio et al., 2024; Rosado et al., 2022; Satoh et al., 2015), warranting further investigation in randomized controlled trials (James et al., submitted; James et al., 2023).

## Author contributions

DM: co-PI, grant writing, conceptualization, investigation, MRI data collection, data curation, formal analysis, and writing, original draft preparation. CEJ: PI, grant writing, funding acquisition, conceptualization, supervision, and writing, review and editing. DK: MRI and behavioral data collection, preliminary preprocessing, and analyses. CJT: psychomotor training program creation, conceptualization, review. MK: input on funding acquisition, conceptualization, and review. GA: clinical recruitment and support, conceptualization, review. ABG: clinical recruitment and support, conceptualization, review. GBF: clinical recruitment and support, conceptualization, review.

## Acknowledgments

We thank the technical staff from the imaging platform at the Brain and Behaviour Laboratory in Geneva (BBL, https://bbl.unige.ch/) and from the Laboratoire de Recherche en Neuroimagerie in Lausanne for their continuous support (https://www.unil.ch/lren/home/menuinst/lreners/technical-support.html). We also thank Hugo Willerval, who designed and delivered the music intervention and made an important contribution to the design of the music practice program. We would also like to thank Aurélie Revol and Angéline Bruneau who provided the psychomotor training courses and greatly participated in its design. We thank Cyrille Stucker and António Manuel Fernandes, the psychologists who coordinated the study and greatly contributed to behavioral and MRI data collection. Finally, we thank our former Master students in Neuroscience, Cécile A.H. Müller and David M. Müller for contributing to MRI and behavioral data collection.

## Data availability

The datasets generated during this study are not publicly available due to patient confidentiality requirements and restrictions imposed by the ethics committees (Commission cantonale d’éthique de la recherche de Genève, CCER, and Commission cantonale d’éthique de la recherche sur l’être humain Vaud, CER-VD), which prohibit data sharing outside Switzerland. Data are available from the corresponding authors on reasonable request subject to ethical approval.

